# An educated guess: how coral reef fish make decisions under uncertainty

**DOI:** 10.1101/2022.08.28.505588

**Authors:** Cait Newport, Adelaide Sibeaux, Guy Wallis, Lucas Wilkins, Theresa Burt de Perera

**Affiliations:** Department of Biology, University of Oxford, Oxford, United Kingdom; Centre for Sensorimotor Performance, School of Human Movement and Nutrition Sciences, University of Queensland, Brisbane, Australia

**Keywords:** Adaptive decision making, learning, fish, cognition, vision, alternative forced-choice test

## Abstract

Making informed decisions is fundamental to intelligent life and is crucial to survival. The problem many organisms face, is how to make optimal decisions in the face of incomplete, unreliable, or conflicting information. In many aquatic environments, fish use visual information to guide key behaviours, but the environment itself can alter or mask the very signals they rely on. Here, we asked how a highly visual species, *Rhinecanthus aculeatus*, responds to a learned discrimination task as signal reliability decreases, and whether probabilistic information gained during previous experience can be incorporated into their decision strategy. Fish were first trained to select a target (dark grey circle) from three distractors (light grey circles). In the first experiment, the target was more likely to appear in one of four possible stimulus positions. In the second experiment, the target appeared in all positions equally. In a series of trials, the difference in brightness between the target and distractors was reduced until all four stimuli were identical. We found that target selection accuracy decreased with decreasing target and distractor disparity. However, in Experiment 1 where the target was more likely to be in one position, fish increasingly selected stimuli in the biased position as target selection accuracy decreased, but not in Experiment 2. These results demonstrate 1) that fish learned more than a simple select/avoid rule based on stimulus brightness, but also integrated information (stimulus position) that could be considered ancillary to the primary task. 2) Fish can learn probability distributions and apply this knowledge as uncertainty increases, ultimately increasing the overall frequency of correct choices. Our results reveal that probabilistic decision rules can be used by fish when visual information is unreliable, indicating a possible mechanism for decision making given the inherent noise in incoming sensory information.

## INTRODUCTION

Vision is an important sensory system guiding a wide range of behaviours in fish (e.g. predation, signalling, spatial cues, etc). Errors in signal processing can lead to non-adaptive behaviour, and ultimately impact the survival and fitness of individuals. However, aquatic visual environments can be incredibly noisy, and sensory pollution can lead to signal masking, distraction or misleading (Dominoni et al., 2020). For example, decreases in water clarity cause scattering and attenuation of light, reducing detection distances and scene contrast. In particular, underwater caustics, caused by the refraction of downwelling light, produce moving bands of light that can interfere with visual signal detection including prey detection (Attwell, Ioannou, Reid, & Herbert-Read, 2021; S. R. Matchette, Cuthill, Cheney, Marshall, & Scott-Samuel, 2020; Samuel R. Matchette, Cuthill, & Scott-Samuel, 2019). While some aquatic species have evolved visual mechanisms to reduce or eliminate sources of noise (e.g. crabs (*Carcinus maenas*) and cuttlefish (*Sepia officinalis*) using polarised light (Venables et al., 2022), others are making decisions in situations where the signal to noise ratio is low and where errors in signal detection or interpretation are expected to be high. Given the importance of accurate signal interpretation in governing behavioural output, it makes sense that animals in dynamic and noisy environments have adaptations that allow them to cope with reduced signal fidelity.

One strategy might be to generalize more across stimuli or to decrease reaction thresholds to important stimuli. For example, any looming stimulus could be treated as a potential predator causing a fish to dart away or hide. While this would minimize the risks of predation, it could also increase the number of times an individual responds to a non-predator stimulus. False alarms would reduce time spent on other behaviours such as foraging and mating, thereby incurring a lesser, but still significant, cost. Indeed, it has been shown that turbid conditions prompt sticklebacks to take longer to leave a refuge area and to consume less food once it was found (Chamberlain & Ioannou, 2019). However, evidence suggests that other species perceive a lower risk in turbid conditions. For example, juvenile chinook salmon (*Oncorhynchus tshawytscha*) will remain close to the bottom on an experimental arena in clear water but will be distributed randomly in turbid conditions (Gregory, 1993; Gregory & Levings, 1996). The introduction of a simulated predator causes the fish to swim deeper in both clear and turbid water, but in turbid conditions they return to their pre-predator behaviour faster (Gregory, 1993). It may be that for these species, weak signals are an indicator of a less immediate threat (Lukas et al., 2021).

An alternative strategy would be to incorporate other sources of information to increase signal reliability (Ernst & Bülthoff, 2004; Kulahci, Dornhaus, & Papaj, 2008; Munoz & Blumstein, 2012). For example, the accumulation of evidence across multiple sensory systems could help replace missing information or enhance signal discrimination. However, this is likely to be task dependent as the information provided by other sensory systems are not always equally informative. For example, weakly electric fish (*Gnathonemus petersii*) can discriminate objects using both their highly evolved electric sense as well as vision (Schumacher, Burt de Perera, Thenert, & von der Emde, 2016), but visual cues are preferred when objects are at a greater distance as they are more reliable than electrosensory cues (Schumacher, Burt de Perera, & von der Emde, 2017). Collective decisions are another mechanism to increase signal detection and identification accuracy for social fish. This is a useful strategy when trying to detect a more global signal such as a predator (Davidson et al., 2021; Ward, Herbert-Read, Sumpter, & Krause, 2011), but not all decisions can be optimised using a group quora, and of course there are costs associated with the increased competition associated with living in large social groups.

In many cases, animals will not have access to additional information, and will instead have to rely on a single sensory system. In this case, how might an animal improve decision-making when faced with signal ambiguity? A Bayesian inference approach incorporates probabilistic knowledge accrued through previous experience and degree of uncertainty into the decision-making process (Gershman, 2015; Ma, 2019). Here we tested if fish could use a similar probabilistic approach to solve a visual discrimination task, and whether their reliance on probabilistic information was related to signal uncertainty. Using an operant learning assay to interrogate animal decision making, we tested whether a species of coral reef fish would integrate previous experience of reward location to optimally weight their choices in the face of variations to the signal strength of their reward stimulus.

## METHODS

### Experimental overview

In two experiments, individual Picasso triggerfish (*Rhinecanthus aculteatus*) were presented with a four-alternative forced-choice task in which there was one dark grey circular target (S+) and three light grey circular distractors (S-) (Fig. 1). Grey stimuli were chosen because a straightforward change in brightness could be used to alter the discrimination difficulty, and because *R. aculeatus* can learn to discriminate them quickly and to a high degree of accuracy. Stimuli were presented in four possible positions: top right (TR), top left (TL), bottom right (BR) and bottom left (BL). The two experiments differed in terms of the target position distribution. In Experiment 1, there was a bias in the target position distribution allowing subjects to learn probability information. In Experiment 2, the target was equally likely to appear in any of the four positions. In Experiment 1, our hypothesis was that fish would fall back on their expectations linked to target location when exposed to conditions of increasing stimulus uncertainty. However, in Experiment 2, in which there was no prior position bias, we did not expect the fish to have a preference for any particular position.

**Figure 1:**
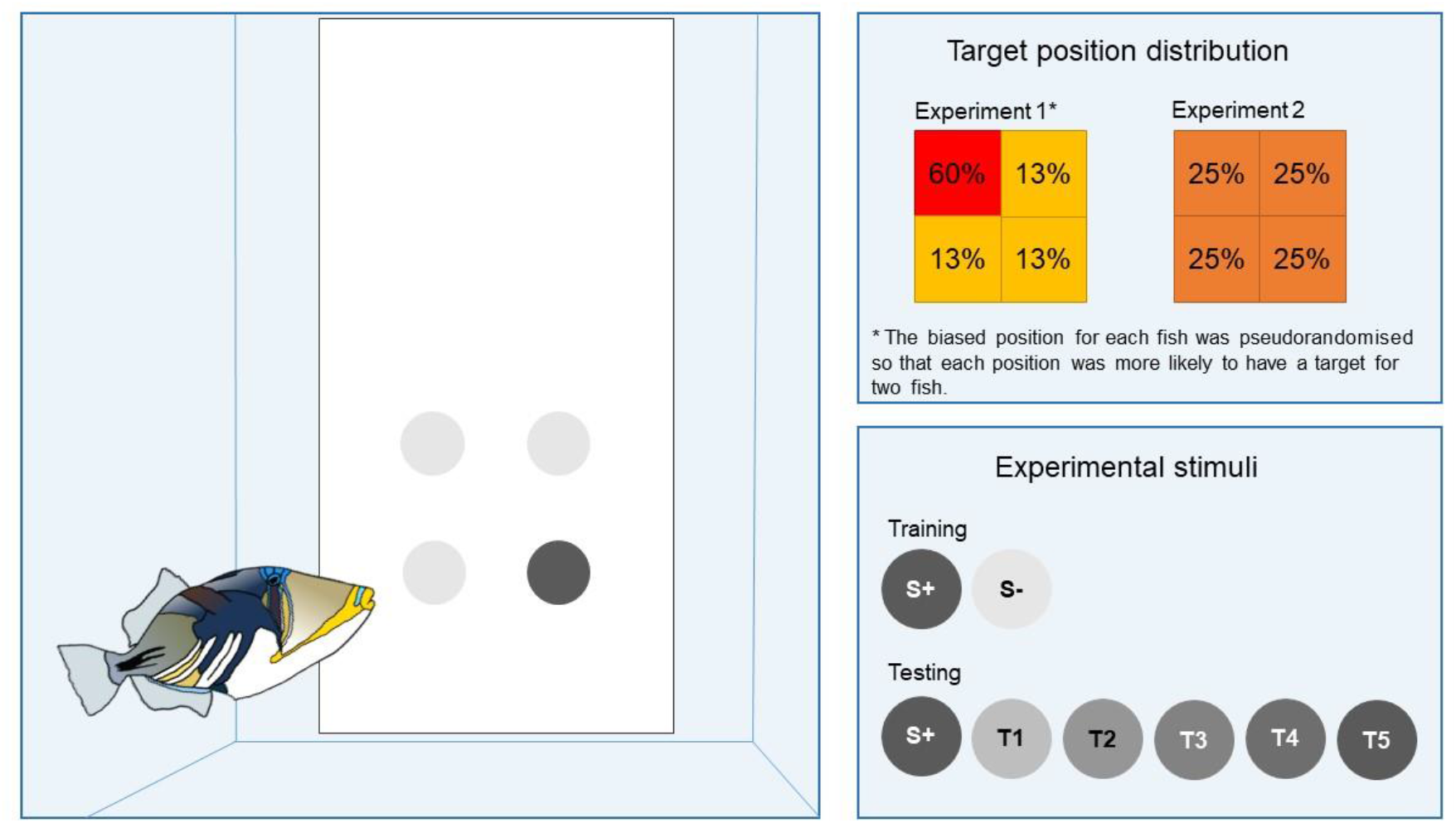
Summary of the experimental protocol. Selection of the target stimulus (S+) was rewarded with food, while the distractor stimuli (S-) were not. The figure is illustrative only and not rendered to scale.

Both experiments were divided into two training steps followed by a testing stage. The first training step taught fish to discriminate stimuli based on colour/brightness and in the second step fish were introduced to the target position distribution. In the second training step, the distribution of target positions was altered such that the likelihood of the target appearing at one location increased relative to the other three (for Experiment 1 only). In Experiment 2, fish still underwent this training step, but the correct stimulus was distributed equally across all four positions. All fish were given 12 sessions (10 trials each) in training step two, regardless of target selection accuracy. Of the 120 trials, in 72 trials (60%) the target was presented in the ‘biased’ position. In the remaining trials, the target appeared 16 times in each of the three other positions. Two fish were trained to each of the four possible positions (TR, TL, BR, BL).

A two-tiered reward system was used in the hope that it would direct the attention of the fish to the secondary rule. When fish correctly selected S+ in the non-biased position, they were given a standard food pellet, but when the fish correctly selected the S+ and it was in the biased position, they were given a larger piece of cut-up bait fish. This reward procedure was used throughout Step 2 and for reinforcement trials in the testing stage. S+ selection accuracy remained high during Step 2 (combined mean frequency for all sessions CI: 95 % ± 1.8 CI) and all fish progressed to the testing stage.

In the testing stage, S-was replaced with distractors that became increasingly difficult to distinguish from the target stimulus. Testing sessions included five reinforcement trials (R) and five probe trials (T1-T5). The reinforcement trials followed the same procedures as those of training Step 2 (S+ and S-) and were used as a reminder of the original task. Correct selections were rewarded, and the two-tiered reward system was maintained in these trials, therefore three of the five reinforcement trials were in the predetermined position. The remaining five trials were probe trials in which the difference in stimulus brightness became increasingly similar and eventually identical. In each testing session, one of each of the five treatments was presented. Probe trials were unrewarded, and the biased position was not maintained. The order of reinforcement and testing trials was pseudorandom so that a particular trial type (R or T) would not appear in more than two consecutive trails. A total of 24 testing sessions were run.

In Experiment 2, the methods used in training and testing were similar to that of Experiment 1, except that fish were not biased to a particular stimulus position in Step 2. In addition, there was a difference in reward value and frequency. For Fish 18K and 17R, the two-tiered reward system was maintained; in 60% of trials (selected pseudo-randomly), one of four positions was preassigned a higher reward value if chosen, with an equal frequency for all four positions. For the other two fish, the reward schedule was simplified so as all correct responses resulted in a single pellet. As in Experiment 1, the probe trials were not rewarded during testing for Fish 18K and 17R. However, Fish 24O and 25P, were rewarded. Fish maintained an S+ selection accuracy of 97% ± 1.5 CI in Step 2.

Under the training scenario, selecting stimuli based on their visual characteristics (i.e. the darkest stimulus) would lead to the highest reward rate (100%). Under the testing condition, as the stimuli become harder to discriminate, the likely reward rate decreases, with the level of discrimination difficulty, to chance (25%). However, by changing decision strategy to include position in their selection criteria, subjects in Experiment 1 could increase their chances of receiving a reward (60%). In this paradigm, two questions emerge. 1) Will the fish use position as a cue, and 2) if they do, will they change their decision strategy in such a way as to maximize their food rewards across all probe types, or will they only rely on position when visual cues are completely unusable?

### Experimental procedure

At the beginning of an experimental session, a white opaque Perspex trapdoor was placed in the front third of the home tank (18cm from the front of the tank) and was used to separate the fish from the remainder of the tank. Once the door was in place, a white stimulus presentation board (40 x 20 cm) was inserted into the final quarter of the tank. There were four possible stimulus positions on the presentation board: top right (TR, 17cm from bottom, 5.5cm from right edge), top left (TL, 17cm from bottom, 5.5cm from left edge), bottom right (BR, 7cm from bottom, 5.5cm from right edge) and bottom left (BL, 7cm from bottom, 5.5cm from left edge).

A trial began with the fish behind the trap door. Once the door was raised, the fish swam into the open area and up to the presentation board and had to select one of four stimuli. If the fish choose the correct stimulus by biting it, they received either a food pellet (Formula One Marine Pellet, Ocean Nutrition, United States) or a piece of fish (Gamma Slice, Tropical Marine Centre, United Kingdom), depending on the experimental trial type. If the fish choose incorrectly, they were not rewarded, the presentation board was removed from the aquarium, and the trial was terminated. After each trial, the fish swam back through the trap door and the door was closed in preparation for the next trial. One session consisting of 10 trials was run each weekday during the experimental period. See Supplementary Fig. 1 for a video example of the testing procedure. In training stage 1, fish were deemed to have learned the task once they achieved >75% accuracy in three consecutive sessions (10 trials/session). Each fish was given as many sessions as required to learn this task. In Experiment 1, all fish (n = 8) completed this stage within 4-11 sessions. In Experiment 2, all fish (n = 4) completed training Step 1 within 3-11 sessions.

### Subjects

A total of 12 fish were used in this experiment; eight individuals were tested in Experiment 1 and another four were tested in Experiment 2. Individuals were purchased from local suppliers. The fish were housed in individual aquaria (35.5 x 60 x 31.5 cm) in a flow through marine aquarium system and fed a daily mixed diet of pellets and fish pieces as part of experiments. All fish had rock and a PVC pipe (length: 15cm; diameter: 10.5cm) to serve as shelter and enrichment, which were moved during experiments to ensure an unobstructed view of the experimental stimuli. Three fish had previously participated in similar experiments using different training techniques, but the remaining fish were naïve (Supplementary Table 1). Prior to this experiment, all fish were at least pre-trained to swim through a trapdoor and to select white discs from a presentation board.

### Ethical note

All fish were cared for according to the code of practice for the care and use of animals for scientific purposes (Local Ethical Review Committee of Oxford University’s Department of Zoology). Fish were held individually to minimise stress associated with territoriality. Experiments were conducted within the home tank of each fish to reduce handling and they were provided with enrichment in the form of shelters, rocks and gravel. Fish were offered live coral prior to experiments however the fish destroyed all soft material within an hour which had an impact on water quality, therefore these were removed and replaced with standard aquarium rocks.

### Stimuli

Stimuli were grey circular discs (2.5 cm diameter) that were printed on photographic paper (Canon matte photo paper), cut out using a hole-punch and laminated. Six stimulus types were made (Fig. 1). The darkest (S+) and brightest (S-) were used in training and reinforcement trials (RGB values: **S+**: 90 90 90, **S-**: 230 230 230). The remaining four stimuli were used as testing stimuli with RGB values: **T1**: 190 190 190, **T2**: 150 150 150, **T3**: 130 130 130, **T4**: 110 110 110). A fifth testing condition (T5) used S+ stimuli for all four options.

### Statistical analysis

For Experiment 1 probe T5, all stimuli presented were identical therefore it is impossible to state whether the fish had made a correct or incorrect decision based on the visual cue. For the sake of analysis, before testing began, one stimulus was randomly assigned as S+ for each trial. As these trials are unrewarded, the designation had no effect on the fish, however selection of this S+ was used to calculate the accuracy of those trials.

The selected stimulus colour and position was recorded for all trials. The results were tallied for each probe type and analysed using a generalised linear mixed model (GLMM) with a binomial distribution (logit-link function) to test whether the selection frequencies of each stimulus type (S+ or position) changed along with probes (using the glmer function in the lme4 package; (Bates, Mächler, Bolker, & Walker, 2015)). Selection frequencies of either S+ (test performed for Experiment 1 and 2) or the biased position (test performed for Experiment 1 only) were included as reponse variables. Probe was included as a fixed effect and Fish ID as a random effect. Finally, we tested if the selection frequency at each position changed along with probes (tests performed for Experiment 1 and 2). For each position, selection frequency was included as the response variable. Probe and Fish ID were added as fixed and random effect respectively.

To determine if the stimulus selection on the non-biased position was significantly different from selection on the biased position (Experiment 1 only) we used GLMM. For each probe we ran a GLMM and included the selection frequencies as the response variable. Position was added as fixed effect and fish ID as a random effect.

The linear relationship between the selection frequency of S+ and the selection of stimuli in either the biased position (Experiment 1) or each of the four possible positions (Experiment 2) was analysed using a linear mixed model (using the lmer function in the lme4 package; (Bates et al., 2015)). Fish ID was included as a random effect. Pseudo-R^2^ was calculated using the Efron pseudo-R square formula where the model residuals are squared, summed, and divided by the total variability in the dependent variable.

For each model (glmer or lmer), we assessed the fit between the model and the data, the residuals normality and the residuals dispersion using simulateResiduals and testDispersion functions (DHARMa package in R (Hartig, 2022)). All data is provided in Supplementary File 2, and all statistical code and outputs are reported in Supplementary File 3.

## RESULTS

### Experiment 1

Fish selected S+ at a frequency higher than chance for R (CI: 95.3 - 97.8%), T1 (CI: 82.4 - 92.4%), T2 (CI: 56.4 – 72.3%), T3 (CI: 38.4 – 55.3%) but not T4 (CI: 24.3 – 39.6%) or T5 (CI: 18.3 – 32.4%). See Supplementary Table 1-2 for individual fish results for Experiments 1 and 2. Moreover, the selection of S+ significantly decreased with decreasing brightness contrast between S+ and the distractor stimuli (all P<0.005, see Supplementary File 2: RS1), except for T4 and T5 trials (P=0.14) where fish selected S+ at the same frequency. While fish selected stimuli in the biased position at a frequency consistent with chance for R (CI: 56.9 – 64.3%; Note that S+ appears in the biased position in 60% of R trials) and T1 (CI: 22.5 – 35.6%), as stimuli became more similar, selection of stimuli in the biased position increased for T2 (CI: 39.1 – 53.7%), T3 (CI: 46.8 – 61.4%), T4 (CI: 48.4 – 62.9%) and T5 (CI: 61.6 – 75.2%). See Supplementary File 2: RS2 for P values. Selection of stimuli in all other positions is significantly different from the biased position except for T1 (Fig. 2a, Supplementary File 2: RS3-RS8). In addition, there was a significant negative relationship between selection frequency of S+ and selection of stimuli in the biased position (Fig. 2b, coefficient of regression β = -0.54, pseudo-R^2^ = 0.76, *p* < 0.001; Supplementary File 2: RS9). Although the fish were unrewarded in the testing conditions, had the training reward schedule held true as the fish likely expected, they would have received a food reward at a rate of at least 50% across probe types, which is much higher than what is predicted by chance (25%).

**Figure 2:**
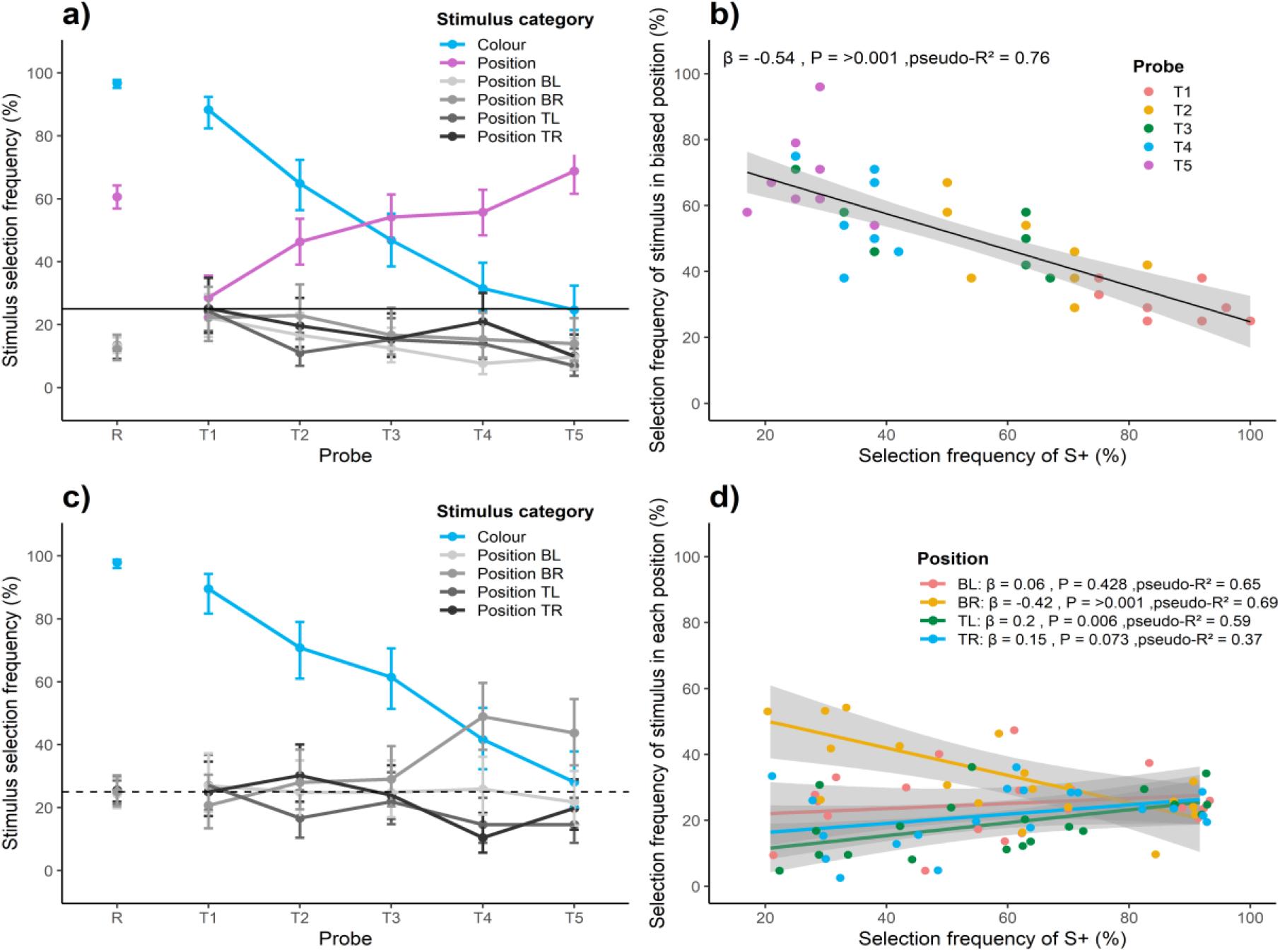
Stimulus selection frequency results of Experiments 1 and 2. In experiment 1, (**a, b**), the target stimulus was 4.5 times more likely to be in one of the four positions when the original conditioned stimuli were presented (S+ and S-). The four possible positions are Top Left (TL), Bottom Left (BL), Bottom Right (BR) and Bottom Left (BL). In experiment 2, (**c**,**d**), target stimulus was presented in all positions equally. **a)** The mean selection frequency of S+ and all stimuli positions for eight fish. The dashed line at 25% indicates a selection frequency consistent with chance. Error bars represent confidence intervals based on a generalized linear mixed effect model. Stimuli used in testing are shown below the figure where the top circles are S+ (not to scale). **b)** A linear regression analysis of S+ selection frequency versus selection of stimuli in the biased position. The shaded grey area indicates confidence intervals. **c)** The mean stimulus selection frequency for four fish, given no position bias. Symbols are the same as those in figure **a. d)** Linear regression analyses of S+ selection frequency versus each of four possible stimuli positions.

### Experiment 2

When the target was equally likely to appear in any of the four positions throughout training, selection accuracy of the target S+ decreased similarly to what was observed in Experiment 1 and was higher than chance for R (CI: 96.2 - 98.9%), T1 (CI: 81.7 - 94.3%), T2 (CI: 60.9 – 79.4%), T3 (CI: 51.4 – 70.3%), T4 (CI: 32.3 – 51.7%) but not T5 (CI: 20.0 – 37.9%). See Supplementary File 2: RS10 for significance of S+ selection frequency between the different probes. The selection frequency of stimuli in either of the four positions was not generally above chance levels across treatments, except in the case of the bottom right position (Position BR) which was significantly higher than chance for T4 (CI: 38.4 – 59.6%) and T5 (CI: 33.5 – 54.5%). See Fig. 2c for further confidence intervals and Supplementary Files RS11 - RS16 for tests on difference in position selection frequencies at each probe. There was a significant relationship between selection frequency of S+ and stimuli in Positions BR (β = -0.418, pseudo-R^2^ = 0.690, *p* <0.001) and TL (β = 0.203, pseudo-R^2^ = 0.587, *p* = 0.006), but not Positions BL (β = 0.056, pseudo-R^2^ = 0.647, *p* = 0.428) or TR (β = 0.153, pseudo-R^2^ = 0.370, *p* = 0.073) (Fig. 2d; Supplementary File 2: RS17).

## DISCUSSION

The experiments in this study demonstrate two important findings: 1) Fish can learn probability distribution information, and 2) they can apply this knowledge in a visual discrimination decision-making task in a manner commensurate with increasing visual uncertainty as to the correct choice. When the fish were presented with stimuli they had learned to a high degree of accuracy, they chose the correct stimulus in almost all trials. However, as their S+ selection error rate increased, the fish relied more on their previous knowledge of the correct response distribution, and increasingly made selections based on position. As a result of this decision strategy, the fish increased their potential for receiving food rewards (Experiment 1). These experiments have thus revealed a powerful mechanism for fish decision making when faced with uncertainty about incoming sensory cues.

Bayesian inference is an application of Bayes’ theorem which describes the integration of prior probability knowledge of a specific outcome with current incoming information, allowing the assessment of probability to be updated as more information becomes available (J. McNamara & Houston, 1980; J. M. McNamara, Green, & Olsson, 2006; Ramírez & Marshall, 2017; Trimmer et al., 2011). It provides a framework for optimal decision making in the face of uncertainty but requires that the decision-maker can learn and remember probabilistic information and alter their expectations based on their current situation. In non-human animals, taxonomically diverse species (e.g. bees, crabs, fish, birds, mammals) have demonstrated selection behaviour consistent with Bayesian inference during a range of behaviours including foraging (Alonso, Alonso, Bautista, & Muñoz-Pulido, 1995; Jay M. Biernaskie & Gegear, 2007; Jay M Biernaskie, Walker, & Gegear, 2009; Foley & Marjoram, 2017; Lima, 1984, 1985; Marshall et al., 2013; Milinski, 1994; Olsson & Noél, 1999; Olsson, Wiktander, Holmgren, & Nilsson, 1999; Valone, 1991, 1992; Valone & Brown, 1989; Valone & Giraldeau, 1993; van Gils, Schenk, Bos, & Piersma, 2003), mate searching (Hunte, Myers, & Doyle, 1985; Luttbeg & Warner, 1999), collective behaviour (Gunji, Murakami, Tomaru, & Basios, 2018), and natal dispersal (Selonen & Hanski, 2010). In this study we have shown that fish can incorporate the statistics of their past experience and their degree of certainty in incoming sensory information, when making decisions. Using this strategy may afford several benefits: 1) fish can use previous experiences to inform current decisions, 2) they can respond flexibly and dynamically to changing situations, and 3) they can make optimized decisions in uncertain conditions.

However, as has been observed in the foraging strategies of other species, the behaviour of the fish was not optimized to receive the most rewards possible. To maximize potential food rewards, subjects should have selected stimuli in the biased position in all trials. Instead, fish appeared to apply ‘probability matching’ whereby the stimulus selection rate matched the learned probability distribution. Probability matching has been previously observed in some, but not all (e.g. Herbranson & Schroeder, 2010; Parducci & Polt, 1958; Wilson & Rollin, 1959) animals including fish (Erika R. Behrend & Bitterman, 1961; E.R. Behrend & Bitterman, 1966; Bullock & Bitterman, 1962; Perez-Escudero & de Polavieja, 2011; Woodard & Bitterman, 1973) and humans (e.g. Gaissmaier & Schooler, 2008; Herbranson & Schroeder, 2010; Rubinstein, 2002). Why it occurs and its evolutionary significance is still debated. One possibility is that it is the result of a cognitive shortcut (Vulkan, 2000), while another possibility is that the randomness of a sequence is not internalized leading to the use of alternative optimizing strategies (e.g. “win-stay, lose-shift”) (Gaissmaier & Schooler, 2008). However, these explanations are typically considered in the context of human decision-making and may not be relevant to all animals, including fish, if such divergent species do not share the same cognitive biases. Both the aforementioned explanations imply that probability matching is an artefact of a cognitive weakness, but it may also provide some benefit to an individual as it allows for sampling of the environment and therefore an opportunity to update priors where conditions may be fluctuating (Pisupati, Chartarifsky-Lynn, Khanal, & Churchland, 2021). Whatever the cause, there is experimental evidence that the application of probability matching may be context dependent, and contingent on incentives, motivation, reinforcement, or understanding of the problem (Bitterman, 1971; Börgers & Sarin, 2000; Erev & Barron, 2005; Rivas, 2013; Shanks, Tunney, & McCarthy, 2002; Vulkan, 2000; Wolford, Newman, Miller, & Wig, 2004). In our Experiment 1, three of the eight fish choose the biased position in T5 at a higher rate than probability matching predicts (fish: 8M = 96%, 9N = 71%, 21P = 79%). Individual differences such as these have been observed in other studies even with very different experimental protocols (e.g. Bitterman, 1971), suggesting that individual variation is a common occurrence. Although the cause or function of probability matching is outside the scope of this study, for the purposes of our results, this behaviour provides clear evidence that fish in Experiment 1 learned the specific probability distributions of the positive stimulus.

Our use of a two-tier reward system in Experiment 1 served the purpose of directing the attention of fish to the biased position, but it also conflated increased reward probability with increased reward magnitude. To our knowledge, there are no experiments specifically testing if fish incorporate reward magnitude into their decisions. However, cleaner wrasse (*Labroides dimidatus)* are well known for their ability to assess the value of a client and refrain from biting high value clients (e.g. large frequent visitors) to ensure those clients return in the future (Wismer et al., 2019). Given the importance of reward magnitude to a wide range of animals, including bees (Gil & De Marco, 2009), pigeons (Rose, Schmidt, Grabemann, & Güntürkün, 2009), and rats (Zoratto, Laviola, & Adriani, 2016), it is likely that reward magnitude also influences fish learning and decision-making. Experiments with pigeons have shown that they learn a colour discrimination task faster when given a larger reward (Rose et al., 2009). Rats are more likely to choose an option with an uncertain outcome if the eventual reward value is significantly higher than the more reliable choice (Zoratto et al., 2016). In our experiment, we do not know for sure whether the bait fish is perceived as a more attractive reward for the fish, or whether there was a novelty effect. Nevertheless, the probability of getting a reward, or the probability of getting a higher reward would ultimately predict the same decision outcome.

The individual variation observed in this, and many studies of fish behaviour are almost certainly a product of signal noise introduced at various stages of the information processing systems (Faisal, Selen, & Wolpert, 2008). It may also be a product of a Bayesian decision strategy. Probability-based choices can lead to more of a gradient of choices than other context-specific decision rules such as “if-then” or “win-stay, lose-shift”, although this depends on the specifics of the rules learned. A probability-based decision rule may explain not only the range of behavioural outcomes observed in animals, but also the variability in their decisions even when faced with what appears to be identical choices (Beck, Ma, Pitkow, Latham, & Pouget, 2012), such as in a controlled laboratory environment. This is because an individual’s priors, their level of uncertainty in a given trial, and whether they will make a choice that favours their expectation of a positive outcome or not, may differ. Our results also show that individual experience, and by extension, memory, plays an important role in fish decision-making. Hard-wired rules presumably have less demand on long-term memory, while the application of probability distributions would require some, and possibly quite a significant capacity for individual memory. How long these types of memories last, and whether or how quickly they can be overwritten by a period of acclimation to new circumstances, are interesting questions for future studies and would be valuable to our understanding of how fish adapt to persistent and stochastic changes in their environment.

Using probability may generalise to a wide range of behaviours (e.g. foraging), but our results may be particularly important to our understanding of sensory signal processing. In our experiments, uncertainty was introduced as the sensory input (visual brightness) approaches the limit of discrimination. Therefore, the results are relevant to our understanding of intermediate signal processing and may extend to the integration of multiple signal types or sources of information. For example, it suggests that fish can make optimized (but perhaps not optimal) decisions about sensory input, even when the signal is weak or ambiguous. In a coral reef fish species that uses visual signals for a range of important behaviours under rapid changes in visual conditions, this capacity may be particularly important for survival. Our results, combined with that of previous studies, provide increasing evidence that Bayesian inference may be generally applicable across taxa and tasks.

## ACKNOWLEDGEMENTS

We thank Hannah Smithson and Brian Rogers for advice on relative reflectance measurements. We thank John McNamara, Alasdair Houston, and Alex Kacelnik for comments on the manuscript. This work was supported by the European Union’s Horizon 2020 research and innovation programme under the Marie Skłodowska-Curie grant agreement [659684], and a Leverhulme Trust Early Career Fellowship. The contents of the article reflect on the authors’ views and not the views of the European Commission.

