## Supplementary Materials for "An educated guess: how coral reef fish make decisions under uncertainty"

### **Electronic Supplementary Information**

**Supplementary Fig. 1.** Video demonstrating the procedure for different trial types during testing.

### **Table 1.** Individual fish stimulus selection frequency

| **Fish ID** | **S+ selection frequency (%)** | | | | | | **Previous experimental experience** |
| --- | --- | --- | --- | --- | --- | --- | --- |
|  | **R (n = 120)** | **T1 (n = 24)** | **T2 (n = 24)** | **T3 (n = 24)** | **T4 (n = 24)** | **T5 (n = 24)** |  |
| *Experiment 1* | | | | | | | |
| 2-B | 100 | 100 | 83 | 67 | 33 | 21 | yes |
| 8-M | 98 | 83 | 63 | 63 | 38 | 29 | yes |
| 9-N | 96 | 75 | 46 | 38 | 38 | 29 | yes |
| 11-D | 98 | 100 | 71 | 58 | 38 | 17 | no |
| 12-L | 94 | 75 | 58 | 33 | 25 | 33 | no |
| 14-M | 95 | 92 | 71 | 33 | 29 | 29 | no |
| 20-Q | 95 | 88 | 75 | 58 | 38 | 17 | no |
| 21-P | 98 | 92 | 50 | 25 | 25 | 25 | no |
| *Experiment 2* | | | | | | | |
| 17-R | 98 | 92 | 63 | 58 | 46 | 21 | no |
| 18-K | 98 | 83 | 71 | 63 | 50 | 29 | no |
| 24-O | 100 | 92 | 88 | 71 | 29 | 33 | no |
| 25-P | 97 | 92 | 63 | 54 | 42 | 29 | no |

### **Table 2.** Individual fish position selection frequency

| **Fish ID** | **Position (BP = Biased Position)** | **Position selection frequency (%)** | | | | | |
| --- | --- | --- | --- | --- | --- | --- | --- |
|  |  | **R (n = 120)** | **T1 (n = 24)** | **T2 (n = 24)** | **T3 (n = 24)** | **T4 (n = 24)** | **T5 (n = 24)** |
| **2-B** | **TL** | 13 | 25 | 17 | 17 | 0 | 8 |
|  | **BL** | 13 | 25 | 21 | 21 | 0 | 8 |
|  | **TR** | 13 | 25 | 21 | 25 | 46 | 17 |
|  | **BR (BP)** | 60 | 25 | 42 | 38 | 54 | 67 |
| **8-M** | **TL** | 14 | 25 | 8 | 21 | 8 | 0 |
|  | **BL** | 12 | 29 | 29 | 12 | 21 | 4 |
|  | **TR** | 12 | 21 | 8 | 8 | 0 | 0 |
|  | **BR (BP)** | 62 | 25 | 54 | 58 | 71 | 96 |
| **9-N** | **TL (BP)** | 60 | 33 | 58 | 67 | 67 | 71 |
|  | **BL** | 15 | 21 | 4 | 4 | 17 | 12 |
|  | **TR** | 13 | 29 | 29 | 25 | 17 | 17 |
|  | **BR** | 12 | 17 | 8 | 4 | 0 | 0 |
| **11-D** | **TL** | 15 | 25 | 13 | 8 | 21 | 4 |
|  | **BL** | 12 | 25 | 12 | 17 | 0 | 8 |
|  | **TR (BP)** | 59 | 25 | 42 | 46 | 50 | 58 |
|  | **BR** | 13 | 25 | 33 | 29 | 29 | 29 |
| **12-L** | **TL (BP)** | 62 | 38 | 38 | 46 | 33 | 58 |
|  | **BL** | 14 | 17 | 17 | 17 | 8 | 17 |
|  | **TR** | 12 | 33 | 38 | 17 | 58 | 21 |
|  | **BR** | 12 | 12 | 8 | 21 | 0 | 4 |
| **14-M** | **TL** | 12 | 25 | 8 | 8 | 8 | 0 |
|  | **BL (BP)** | 59 | 25 | 42 | 58 | 42 | 58 |
|  | **TR** | 13 | 25 | 21 | 17 | 8 | 8 |
|  | **BR** | 15 | 25 | 29 | 17 | 42 | 33 |
| **20-Q** | **TL** | 12 | 25 | 17 | 21 | 29 | 13 |
|  | **BL** | 13 | 17 | 17 | 4 | 0 | 8 |
|  | **TR (BP)** | 63 | 33 | 29 | 50 | 50 | 63 |
|  | **BR** | 13 | 25 | 38 | 25 | 21 | 17 |
| **21-P** | **TL** | 14 | 21 | 4 | 17 | 17 | 17 |
|  | **BL (BP)** | 60 | 25 | 67 | 71 | 79 | 79 |
|  | **TR** | 14 | 21 | 4 | 4 | 0 | 0 |
|  | **BR** | 12 | 33 | 25 | 8 | 4 | 4 |
| *Experiment 2* | | | | | | | |
| **17-R** | **TL** | 24 | 25 | 13 | 13 | 8 | 4 |
|  | **BL** | 25 | 25 | 17 | 13 | 4 | 8 |
|  | **TR** | 25 | 29 | 38 | 29 | 17 | 33 |
|  | **LR** | 26 | 21 | 33 | 46 | 71 | 54 |
| **18-K** | **TL** | 25 | 29 | 17 | 21 | 25 | 17 |
|  | **BL** | 24 | 38 | 29 | 46 | 42 | 25 |
|  | **TR** | 26 | 25 | 29 | 17 | 4 | 17 |
|  | **BR** | 24 | 8 | 25 | 17 | 29 | 42 |
| **24-O** | **TL** | 25 | 21 | 25 | 17 | 8 | 8 |
|  | **BL** | 25 | 25 | 25 | 25 | 29 | 33 |
|  | **TR** | 26 | 21 | 25 | 29 | 8 | 4 |
|  | **BR** | 24 | 33 | 25 | 29 | 54 | 54 |
| **25-P** | **TL** | 23 | 33 | 13 | 38 | 17 | 29 |
|  | **BL** | 25 | 21 | 29 | 17 | 29 | 21 |
|  | **TR** | 27 | 21 | 29 | 21 | 13 | 25 |
|  | **BR** | 26 | 25 | 29 | 25 | 42 | 25 |
